# PGR5 promotes energy-dependent non-photochemical quenching to enable efficient C_4_ photosynthesis under fluctuating light

**DOI:** 10.1101/2024.09.15.613160

**Authors:** Russell Woodford, Jacinta Watkins, Marten Moore, Samuel J. Nix, Suyan Yee, Kai Xun Chan, Barry Pogson, Susanne von Caemmerer, Robert T. Furbank, Maria Ermakova

## Abstract

PROTON GRADIENT REGULATION 5 (PGR5) is essential for generating proton motive force across thylakoid membranes in C_3_ plants and supporting photoprotection under fluctuating light conditions. It is proposed that this function is achieved by regulating cyclic electron flow around Photosystem I. During the evolutionary transition from C_3_ to C_4_ photosynthesis, the leaf abundance of PGR5 has increased, coinciding with a rise in the cyclic electron flow rate. To investigate the contribution of PGR5 to photoprotection in C_4_ photosynthesis, we generated model C_4_ monocot *Setaria viridis* with null *pgr5* alleles. We show that plants lacking PGR5 struggle to establish proton motive force and energy-dependent non-photochemical quenching (qE) at higher irradiances during instantaneous measurements. This leads to a progressive decline in maximum Photosystem I activity when leaves are exposed to repeated cycles of high irradiance. Additionally, plants without PGR5 exhibit severely reduced growth and photosynthesis compared to wild type plants when grown under fluctuating daylight but not under constant daylight. In the absence of PGR5, a slower-relaxing, zeaxanthin-dependent form of non-photochemical quenching supports growth under fluctuating light, albeit at the cost of reduced photochemical efficiency and assimilation rate. Our findings underscore the role of PGR5 in enabling efficient C_4_ photosynthesis under fluctuating light by establishing proton motive force for the rapid upregulation of qE and preventing photodamage to the electron transport machinery. This study highlights the importance of various non-photochemical quenching mechanisms for C_4_ photosynthesis and emphasises the role of PGR5 in the evolution of C_4_ plants.

## Introduction

C_4_ plants have evolved to outperform C_3_ plants in warm, droughted and nitrogen limited climates by implementing a metabolic C_4_ cycle which operates as a biochemical CO_2_ concentrating mechanism (CCM) across mesophyll (Mes) and bundle sheath (BS) cells (Sage and Kubien, 2007, Ghannoum et al., 2011, Sage et al., 2012). The C_4_ cycle begins in Mes cells where carbonic anhydrase converts CO_2_ to bicarbonate, which is then used by phosphoenolpyruvate (PEP) carboxylase to produce oxaloacetate (Ermakova et al., 2024). Malate dehydrogenase then reduces oxaloacetate to malate, which diffuses to BS cells. In the most studied and agriculturally relevant subtype of C_4_ photosynthesis, NADP^+^-dependent malic enzyme (NADP-ME) decarboxylates malate in BS chloroplasts to provide CO_2_ and form pyruvate (von Caemmerer and Furbank, 2003, Hatch et al., 1975). Pyruvate returns to Mes cells where is it regenerated into PEP by pyruvate, phosphate dikinase using 2 mol of ATP, concluding the C_4_ cycle. Operation of the C_4_ cycle builds up CO_2_ partial pressure in BS chloroplasts where ribulose-1,5-bisphosphate carboxylase-oxygenase (Rubisco) resides, allowing the enzyme to operate at maximum carboxylation efficiency. This minimises the Rubisco oxygenation reaction and enables the improved growth of C_4_ plants in conditions where productivity of C_3_ plants is significantly reduced due to photorespiration (von Caemmerer and Furbank, 2003, Furbank, 2011, Hatch, 1987).

Due to the benefits of C_4_ photosynthesis, there is significant interest in understanding its key components and exploring how they can be utilised to enhance the productivity of both C_3_ and C_4_ crops (Croce et al., 2024, Ermakova et al., 2021a). However, doing this requires an intricate understanding of chloroplast electron transport pathways which power metabolic reactions of Mes and BS cells and are known to limit the CO_2_ assimilation rate of C_4_ plants at ambient CO_2_ (Ermakova et al., 2023). This is because the operation of the C_4_ cycle requires ATP and, for each molecule of CO_2_ fixed, C_4_ plants need an ATP/NADPH ratio of at least 5/2, compared to the C_3_ plants’ ratio of at least 3/2 (Ermakova et al., 2020). Moreover, due to the increased ATP demand, C_4_ plants are more widespread in areas with high irradiation (Sage et al., 1999), likely increasing their need for efficient photoprotection mechanisms to safeguard electron transport machinery.

Chloroplast electron transport chains of Mes and BS cells have developed separate functions to cater for the increased ATP demands of C_4_ photosynthesis (Munekage and Taniguchi, 2016, Ermakova et al., 2024). In Mes, linear electron flow (LEF) is prevalent in which Photosystem II (PSII) uses light energy to split water and obtain electrons which pass through the Cytochrome *b*_6_*f* (Cyt*b*_6_*f*) complex to Photosystem I (PSI) and ferredoxin (Fd), producing NADPH. During this process Cyt*b*_6_*f* translocates protons across the thylakoid membrane to generate proton motive force (*pmf*) which ATP synthase uses to produce ATP. Since LEF produces ATP and NADPH at a fixed ratio of ∼2.6/2, to increase ATP/NADPH, plants also operate cyclic electron flow (CEF) (Kramer and Evans, 2011). CEF returns electrons from the reducing side of PSI to Cyt*b*_6_*f,* allowing generation of additional *pmf* without net NADPH production (Yamori and Shikanai, 2016). The CEF/LEF ratio in Mes cells of the model C_4_ monocot *Setaria viridis* is predicted to be about 0.1 whilst in BS cells it is confirmed to be 5 (Bellasio and Ermakova, 2022, Ermakova et al., 2024). Several CEF pathways have been proposed in higher plants: a chloroplast NADH dehydrogenase-like complex (NDH)–mediated pathway, a PROTON GRADIENT REGULATION 5 (PGR5)/PGR5-LIKE PHOTOSYNTHETIC PHENOTYPE 1 (PGRL1)–dependent pathway, and a direct PSI-Fd-Cyt*b*_6_*f* pathway (Yamori and Shikanai, 2016, Strand et al., 2017, Ruhle et al., 2021, Malone et al., 2021). As the abundance of NDH, PGR5 and PGRL1 has increased during the evolutionary transition from C_3_ to C_4_ photosynthesis, it is considered that C_4_ plants upregulate these CEF pathways to accommodate their increased ATP requirements (Ishikawa et al., 2016, Munekage and Taniguchi, 2016, Nakamura et al., 2013). Consistent with this, NDH-mediated CEF is the major electron transport route in BS cells of *S. viridis* facilitating generation of ATP for the Calvin cycle (Ermakova et al., 2024). However, the contribution of PGR5/PGRL1 to CEF and ATP production in C_4_ photosynthesis is still unclear.

While CEF is essential for accommodating the ATP demands of photosynthesis, it also has an important function in photoprotection. Underutilisation of NADPH by downstream metabolic reactions quickly results in the over-reduction of the electron transport chain and leads to the formation of triplet chlorophyll (Chl) causing the generation of reactive oxygen species (ROS) and damage to photosynthetic machinery (Murchie and Niyogi, 2011, Lima-Melo et al., 2021, Das and Roychoudhury, 2014). This is particularly prevalent under fluctuating light environments in which rapid changes in irradiance cause frequent imbalances between the production and consumption of ATP and NADPH (Allahverdiyeva et al., 2015, Long et al., 2022, Kromdijk et al., 2016). In reduced conditions, O_2_ accepting electrons from PSI results in the formation of the highly reactive superoxide radical (O_2_^●-^). Plants implement superoxide dismutases (SODs) to protonate O_2_^●-^ and form the less-reactive signalling molecule H O, which serves as an indicator of oxidative stress levels in a signalling pathway between chloroplasts and the nucleus (Chan et al., 2016). Overaccumulation of ROS, however, may result in severe oxidative stress and damage to photosynthetic complexes. To overcome this, plants have developed a suite of photoprotective mechanisms which help to prevent and mitigate oxidative stress. There are multiple auxiliary electron pathways, including CEF, which dynamically adjust the ATP/NADPH supply to match metabolic demands (Walker et al., 2020). Additionally, non-photochemical quenching (NPQ) processes can dissipate a portion of light energy absorbed in the PSII antennae as heat before it reaches the reaction centres, thereby lowering overall electron transport rate (Ruban, 2016, Muller et al., 2001).

To accommodate for both short- and long-term increases in irradiance, plants implement different forms of NPQ. Energy-dependent quenching (qE), the most rapidly-relaxing form of NPQ, is driven by changes in pH of the thylakoid lumen due to the build-up of *pmf*. When the lumen pH drops below 5, protonation of the PSII subunit S (PsbS) triggers conformational changes within the light-harvesting complex II (LHCII) promoting dissipation of absorbed light energy. Concomitantly, conversion of violaxanthin to zeaxanthin de-excites singlet Chl and prevents triplet Chl formation (Jahns and Holzwarth, 2012, Muller et al., 2001). On an intermediate timescale, zeaxanthin-dependent quenching (qZ) acts independently of PsbS and *pmf* and instead involves a slowly interconverting pool of violaxanthin and zeaxanthin promoting the aggregation of LHCII subunits and modulating the structure of thylakoid membranes rather than actively quenching absorbed light energy (Nilkens et al., 2010, Malnoë, 2018, Jahns and Holzwarth, 2012). Under prolonged light stress, plants initiate photoinhibitory quenching (qI), a slowly responding and often irreversible form of NPQ in which the D1 subunit of PSII is inactivated and degraded to prevent PSII activity (Muller et al., 2001, Jahns and Holzwarth, 2012).

PGR5/PGRL1 pathway plays a central role in photoprotection of C_3_ plants. PGR5 is a soluble, stromal protein which can form a dimer-dimer complex with the transmembrane PGRL1 (Hertle et al., 2013). While PGRL1 helps maintain PGR5 stability and activity, the PGR5 protein is more likely to be actively involved in regulating electron transport (Ruhle et al., 2021). Compared to wild type (WT) plants, C_3_ plants lacking PGR5 have lower *pmf* and NPQ as well as reduced availability of PSI electron acceptors and impaired electron transport rates at high irradiances (Suorsa et al., 2012, Suorsa et al., 2016, Suorsa et al., 2013, Yamamoto and Shikanai, 2019, Shikanai et al., 1999, Munekage et al., 2002). While having normal growth under moderate constant daylight, C_3_ *pgr5* mutants are unable to grow under fluctuating daylight when the rapid induction and relaxation of qE is essential for photoprotection (Tikkanen et al., 2010, Suorsa et al., 2012, Long et al., 2022, Niu et al., 2023). Therefore, the major function of PGR5/PGRL1 in C_3_ plants is prompt upregulation of *pmf* to trigger qE, but the exact molecular mechanism of PGR5/PGRL1 is still unclear. Moreover, a role of PGR5 in preventing photosynthetic oscillations has provided evidence for the involvement of the pathway in adjusting ATP/NADPH in C_3_ photosynthesis (Degen et al., 2024, Shikanai, 2024).

Due to the potential involvement in both photoprotection and ATP generation, the role of PGR5/PGRL1 pathway in C_4_ photosynthesis is of great interest. A limited number of studies have investigated the function of PGR5 in C_4_ species, predominantly in the NADP-ME model dicot *Flaveria bidentis. F. bidentis* with *pgr5* knock-down has lower NPQ at irradiances above 1000 μmol m^−2^ s^−1^ and a 20% lower net CO_2_ assimilation at 2000 μmol m^−2^ s^−1^, compared to WT, but shows no difference in biomass or Chl content when grown under constant medium (250 μmol m^−2^ s^−1^) or high irradiance (1000 μmol m^−2^ s^−1^) (Ogawa et al., 2023). Comparatively, overexpression of PGR5 in *F. bidentis* results in a lower PSI acceptor side limitation upon a shift from low to high irradiance, compared to WT, but *pmf,* NPQ, quantum yield of PSII, Chl content, CO_2_ assimilation rates and growth are unaltered (Tazoe et al., 2020). These initial studies suggest that PGR5 facilitates CEF and helps establish NPQ in C_4_ plants. However, a conclusive evidence for the role of PGR5 in qE during C_4_ photosynthesis is still lacking, particularly under fluctuating light conditions and in C_4_ monocots, which are more representative models for major C_4_ crops like maize, sorghum and sugarcane. To address this knowledge gap, we used CRISPR/Cas9 to generate *S. viridis* lacking PGR5 to elucidate the importance of PGR5 to C_4_ photosynthesis.

## Results

### Growth and photosynthesis of PGR5-deficient *S. viridis* under constant and fluctuating daylight

To create *S. viridis* lines with null *pgr5* alleles, Cas9 was targeted to the first exon of *pgr5*. Four new *pgr5* alleles with various single nucleotide indels were obtained, *pgr5-1*, *pgr5-2*, *pgr5-3* and *pgr5-4*, all resulting in a premature stop codon within the chloroplast transit peptide of PGR5 (Fig. S1). Plants with homozygous null *pgr5* alleles were selected and confirmed to lack the PGR5 protein through immunoblotting of leaf protein extracts with PGR5 antibody (Fig. S1). When grown under a constant daylight of 380 µmol m^−2^ s^−1^, plants lacking PGR5 were morphologically similar to WT, had the same leaf Chl content and accumulated the same aboveground biomass at harvest (45 days), apart from *pgr5-4* which had slightly reduced biomass (Fig. 1 a-c). Electron fluxes in leaves were analysed at ambient conditions with MultispeQ. No differences in the effective quantum yield of PSII (φ_II_), the yield of non-regulated non-photochemical reactions (φ_NO_) or the yield of NPQ (φ_NPQ_) were found between plants lacking PGR5 and WT when grown under constant daylight (Fig. 1 d-f). In sharp contrast, under fluctuating daylight (repeated cycles of 5 min at 250 µmol m^−2^ s^−1^ and 1 min at 1000 µmol m^−2^ s^−1^), plants lacking PGR5 were much smaller than WT plants, had on average 50% less Chl and accumulated 89% less biomass. Moreover, during the low light phases of the fluctuating daylight regime, φ_II_ was 42% lower in plants lacking PGR5 while φ_NPQ_ was increased 2.4-fold, compared to WT (Fig. 1 d-f). Interestingly, when grown at high constant daylight of 1000 μmol m^−2^ s^−1^, WT and PGR5-deficient plants did not show differences in growth or photosynthesis (Fig. S2).

**Fig. 1.**
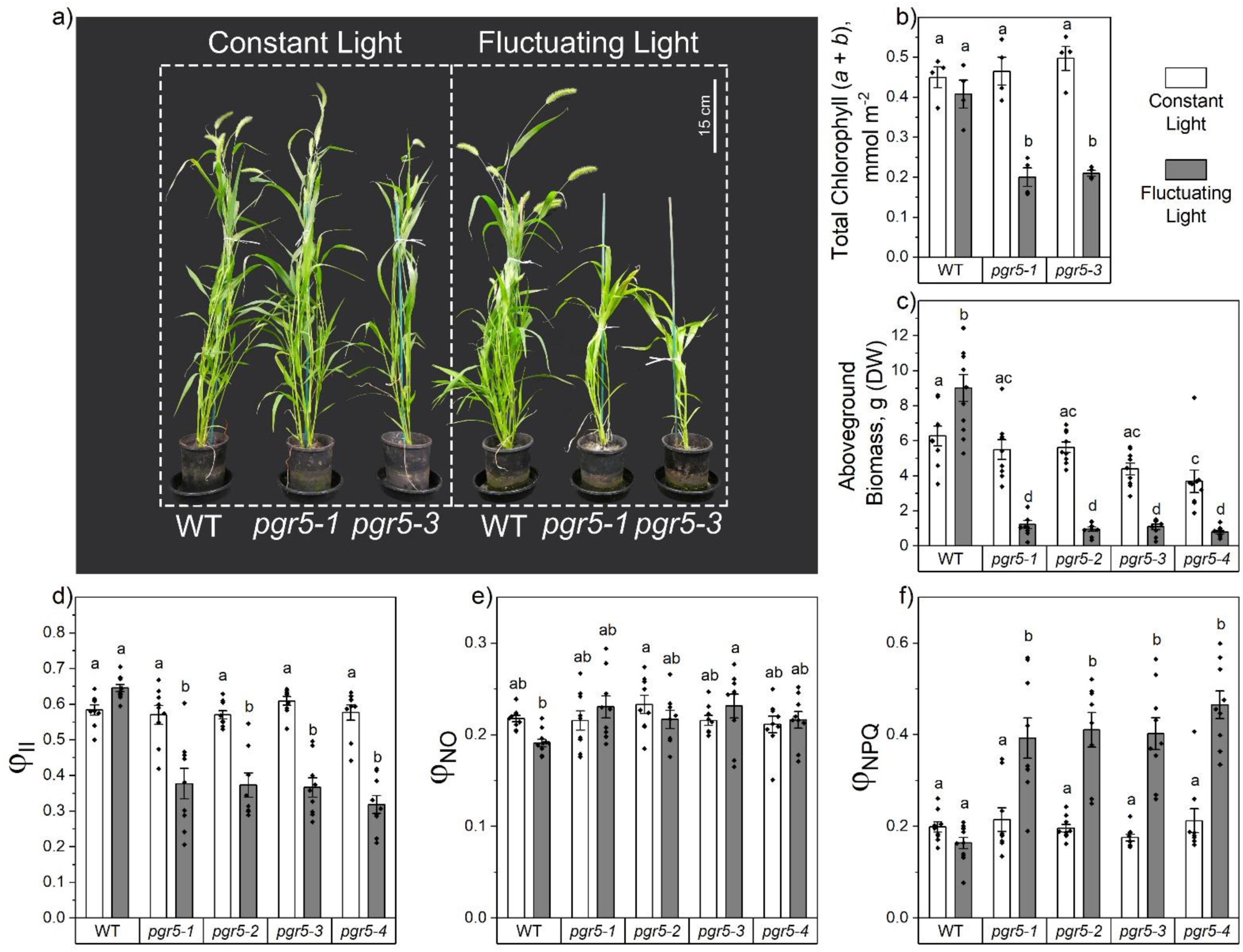
Growth and photosynthesis of wild type (WT) *Setaria viridis* and plants with null *pgr5* alleles grown under constant or fluctuating daylight. (**a**) Growth phenotype of plants 45 days after germination. (**b**) Total leaf Chlorophyll (*a* + *b*) content. (**c**) Total aboveground biomass at harvest; DW, dry weight. (**d - f**) Leaf photosynthesis parameters analysed with MultispeQ at ambient light (380 µmol m^−2^ s^−1^ for constant light, 250 µmol m^−2^ s^−1^ for fluctuating light at low light phase): the effective quantum yield of PSII (φ_II_), the yield of non-regulated non-photochemical reactions (φ_NO_) and the yield of non-photochemical quenching (φ_NPQ_). Mean ± SE, *n* = 4 biological replicates for (b) and *n =* 8-11 biological replicates for (c-f). Letters indicate significant differences between groups (2-Way ANOVA with Tukey’s post hoc test at *P* < 0.05). No differences were found between plants with null *pgr5* alleles.

Gas-exchange analysis revealed drastically different CO_2_ assimilation capacity of PGR5-deficient plants depending on the growth light regime. Net CO_2_ assimilation rates measured at 500 µmol m^−2^ s^−1^ and different CO_2_ partial pressures were similar between plants lacking PGR5 and WT grown at constant daylight and WT grown at fluctuating daylight (Fig. 2a). In contrast, assimilation rates of PGR5-lacking plants grown at fluctuating daylight were severely reduced at all CO_2_ partial pressures, compared to WT. While stomatal conductance was similar between WT and PGR5-deficient plants grown at constant daylight, PGR5-deficient plants grown under fluctuating daylight had greatly impaired stomatal conductance, compared to WT, across all CO_2_ partial pressures (Fig. 2b). We also analysed net CO_2_ assimilation rate at ambient CO_2_ (400 µmol mol^−1^) under gradually increasing and then gradually decreasing irradiances (Fig. 2c). While WT and PGR5-deficient plants grown under constant daylight showed similar CO_2_ assimilation rates as irradiance increased, upon the return from high to low irradiances, plants lacking PGR5 had lower assimilation rates. Fitting of the light response curves showed that the quantum yield of CO_2_ assimilation significantly decreased after the high light exposure in the absence of PGR5 (Φ vs Φ*, Table S1). PGR5-deficient plants grown under fluctuating daylight again showed drastically decreased CO_2_ assimilation rates which further lowered upon the return from high to low irradiances, compared to WT. The dark respiration rate (*R*_d_) and light compensation point (LC) did not differ between WT and PGR5-deficient plants grown under constant daylight while PGR5-deficient plants grown under fluctuating daylight had lower *R*_d_ and higher LC, compared to WT (Table S1).

**Fig. 2.**
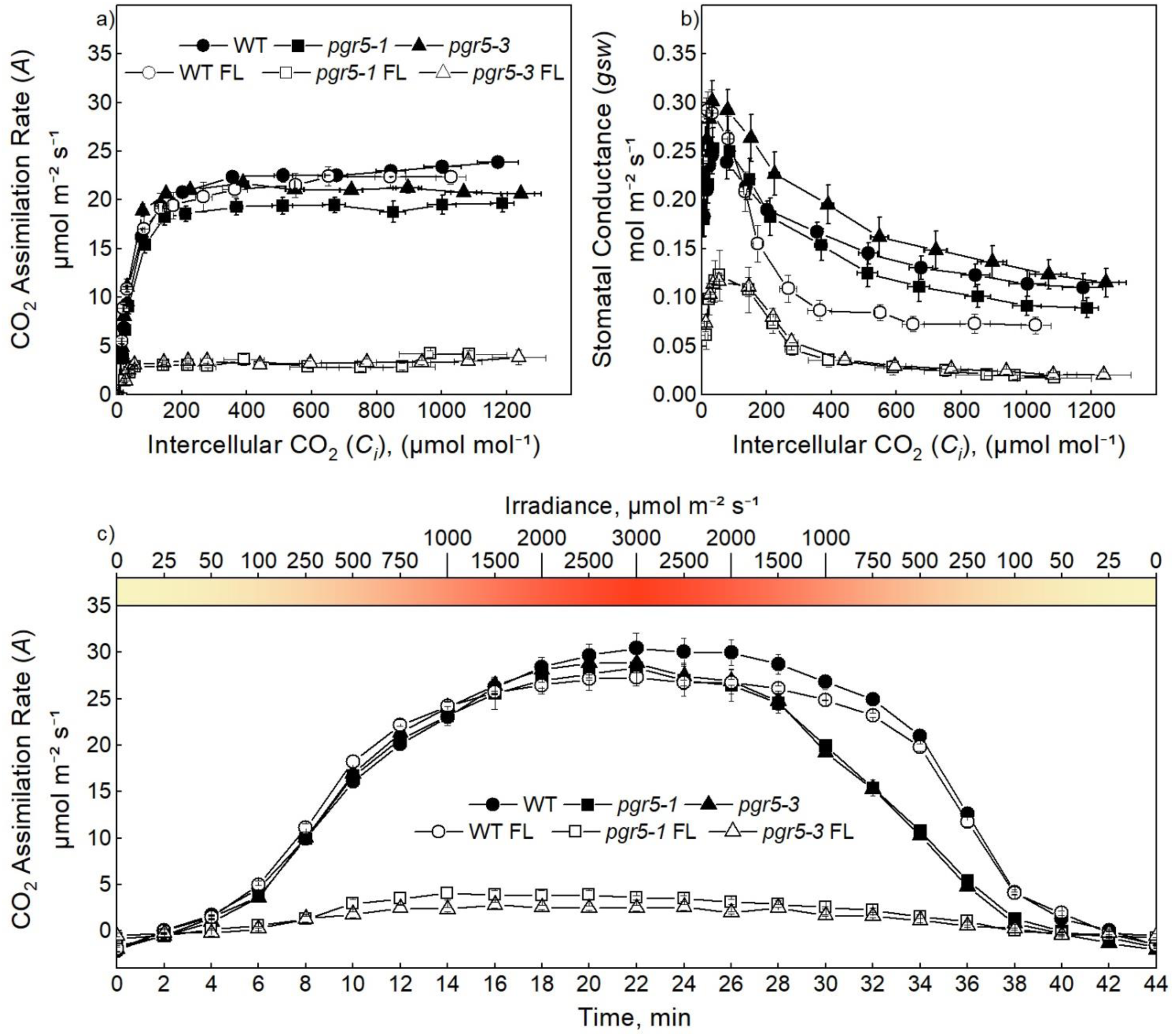
Gas-exchange analysis of wild type (WT) *Setaria viridis* and plants lacking PGR5 (*pgr5-1* and *pgr5-3*) grown under constant (black symbols) or fluctuating daylight (FL; white symbols). (**a**) Response of net CO_2_ assimilation rate (*A*) to the intercellular CO_2_ partial pressure (*C*_i_) at 500 µmol m^−2^ s^−1^. *A* was significantly decreased in FL *pgr5-1* and *pgr5-3* plants compared to FL WT at *C*_i_ above 20 μmol mol^−1^ (1-Way ANOVA with Tukey’s post hoc test at *P* < 0.05). (**b**) Response of stomatal conductance (*gsw*) to the intercellular CO_2_ partial pressure (*C*_i_) at 500 µmol m^−2^ s^−1^. (**c**) Light curves of *A* at ambient CO_2_ (400 μmol mol^−1^), during gradually increasing and then gradually decreasing irradiance. *n* = 4-5 biological replicates. Photosynthetic parameters were fit from the light curves according to von Caemmerer (2021) (Table S1).

Next, we studied activities of PSII and PSI during rapid changes in irradiances through simultaneous measurements of Chl fluorescence and P700 redox spectroscopy (Fig. 3). When grown under constant daylight, plants lacking PGR5 had a significantly lower φ_II_ and higher φ_NO_ across all irradiances, compared to WT, as well as a significantly lower φ_NPQ_ at irradiances above 700 μmol m^−2^ s^−1^ (Fig. 3a-c). Plants lacking PGR5 also had a significantly lower quantum yield of PSI (φ_I_), compared to WT, across all irradiances (Fig. 3g). Furthermore, as light intensity increased, PGR5-deficient plants showed a significant increase in the non-photochemical yield of PSI due to acceptor side limitation (φ_NA_) and a concomitant decrease in the non-photochemical yield of PSI due to donor side limitation (φ_ND_), compared to WT (Fig. 3h, i). Light responses of PSI and PSII yields in plants grown under fluctuating daylight were similar to those under constant daylight but with more extreme differences between PGR5-deficient and WT plants. φ_II_ was drastically decreased while φ_NO_ was drastically increased in plants lacking PGR5, compared to WT, at all irradiances (Fig. 3d, e). φ_I_ was also drastically decreased in PGR5-deficient plants, compared to WT, while φ_ND_ was decreased at irradiances above 700 μmol m^−2^ s^−1^ (Fig. 3j, l).

**Fig. 3.**
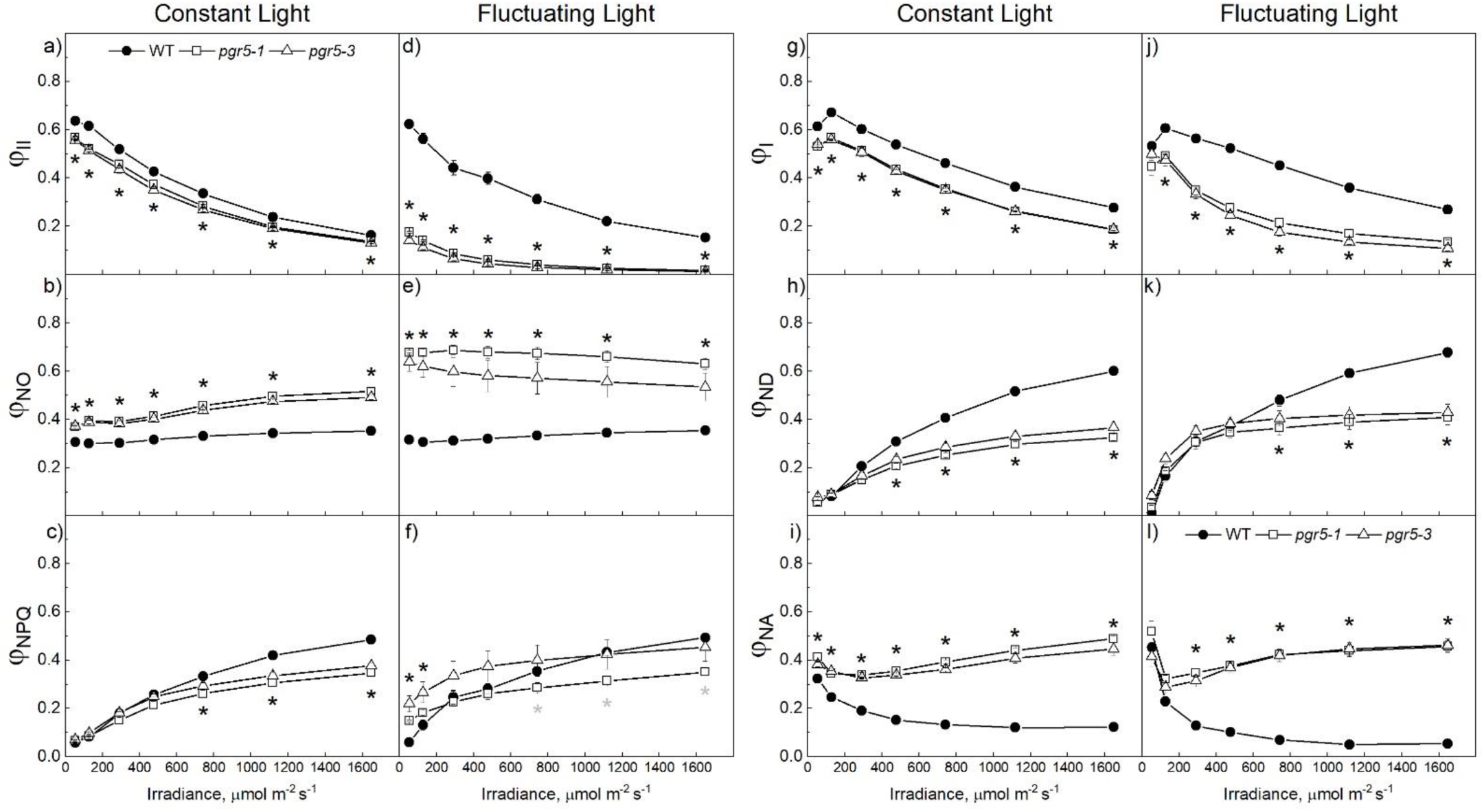
Light response of photochemical yields of Photosystem II (PSII, **a** – **f**) and Photosystem I (PSI, **g** – **l**) in wild type (WT) *Setaria viridis* and plants lacking PGR5 (*pgr5-1* and *pgr5-3*) grown under constant or fluctuating daylight. (**a**, **d**) The effective quantum yield of PSII (φ_II_). (**b**, **e**) The yield of non-regulated non-photochemical reactions (φ_NO_). (**c**, **f**) The yield of non-photochemical quenching (φ_NPQ_). (**g**, **j**) The quantum yield of PSI (φ_I_). (**h**, **k**) The non-photochemical yield of PSI due to acceptor side limitation (φ_NA_). (**i**, **l**) The non-photochemical yield of PSI due to donor side limitation (φ_ND_). Mean ± SE, *n* = 4-5 biological replicates. Black asterisks indicate significant differences between both PGR5-deficient lines and WT plants. Gray asterisks indicate significant differences between *pgr5-1* and WT plants. (1-Way ANOVA with Tukey’s post hoc test at *P* < 0.05).

### Protein, ROS and transcript analyses

Immunoblotting of photosynthetic proteins representing major thylakoid complexes revealed more drastic differences between PGR5-deficient and WT plants grown under fluctuating daylight compared to constant daylight conditions (Fig. 4a-d). Under constant daylight, relative abundances of the NdhH subunit of NDH and the PsaB subunit of PSI were decreased by ∼25% in leaves of PGR5-deficients plants, compared to WT. Abundances of the RbcL subunit of Rubisco, the D1 subunit of PSII, the Rieske subunit of Cyt*b*_6_*f*, the AtpB subunit of ATP synthase and PsbS were unaltered. In contrast, PGR5-deficient plants grown under a fluctuating daylight demonstrated a significant, ∼50% decrease in relative abundances of NDH, PSI and Cyt*b*_6_*f* subunits, compared to WT. Moreover, there was a strong (but significant only in one line) increase in the abundance of PsbS in PGR5-deficient plants, compared to WT.

**Fig. 4.**
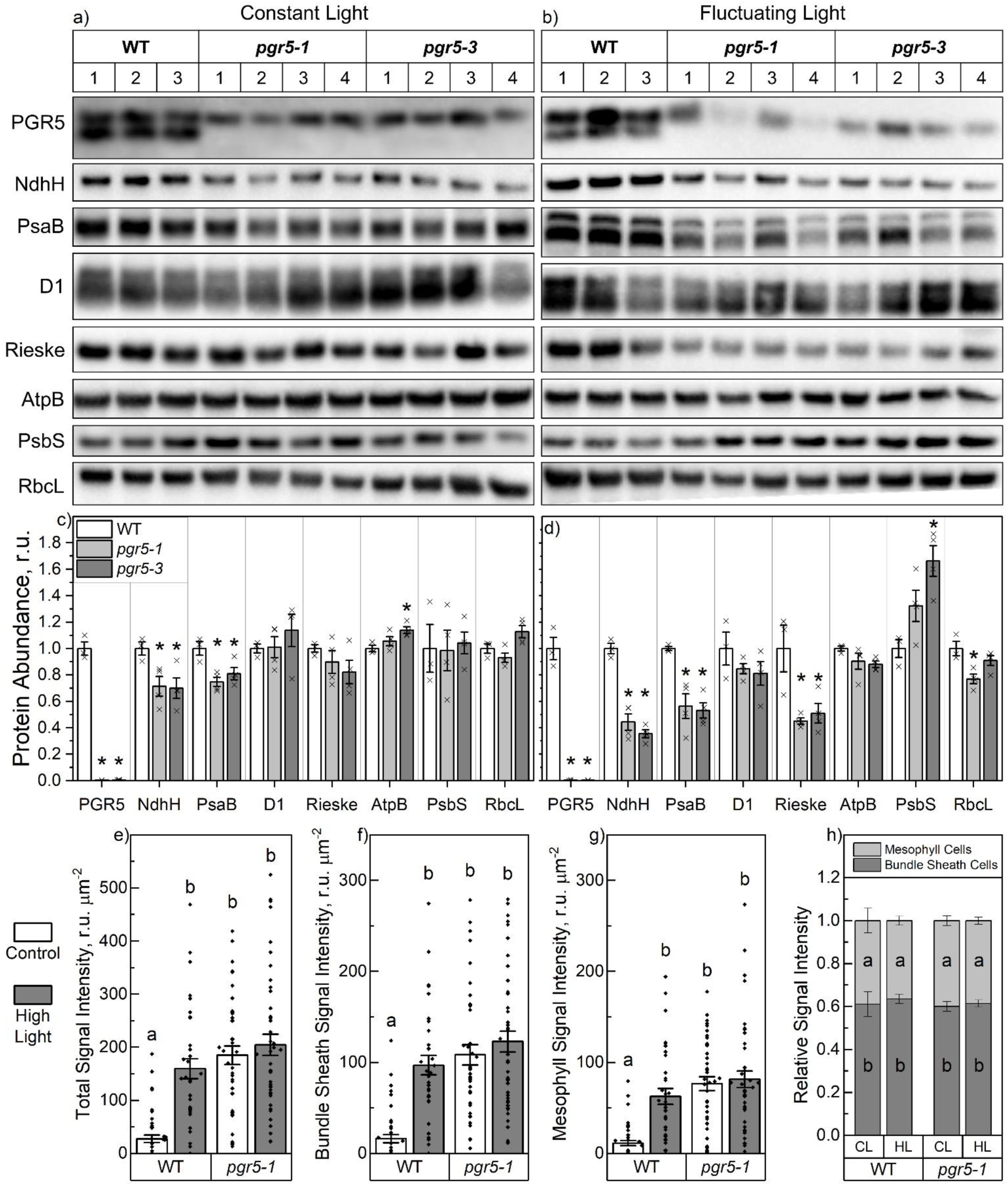
Abundance of photosynthetic proteins and reactive oxygen species (ROS) in wild type (WT) *Setaria viridis* and plants lacking PGR5 (*pgr5-1* and *pgr5-3*). (**a, b**) Immunodetection of PGR5, NdhH (NDH), PsaB (Photosystem I), D1 (Photosystem II), Rieske (Cytochrome *b*_6_*f*), AtpB (ATP synthase), PsbS and RbcL (the large subunit of Rubisco) in leaf protein samples isolated from plants grown under constant or fluctuating light and loaded on leaf area basis. Three to four biological replicates (1, 2, 3, 4) were loaded for each group. (**c, d**) Relative quantification of protein abundance per leaf area. Mean ± SE with three-four replicates shown. Each protein has its own relative scale. Asterisks indicate statistically significant differences between PGR5-deficient and WT plants (1-Way ANOVA with Tukey’s post hoc test at *P* < 0.05). (**e - h**) 2′,7′-dichlorofluorescein diacetate (H_2_DCFDA) fluorescence signal from leaves, bundle sheath and mesophyll cells of constant daylight-grown plants at ambient light (Control, CL) or after 30-min exposure to 2000 µmol m^−2^ s^−1^ (high light, HL). Mean ± SE. Six leaf sections were analysed for each condition, with eight Mes:BS wreath units quantified for each section. Letters indicate significant differences between groups (2-Way ANOVA with Tukey’s post hoc test at *P* < 0.05).

Although the growth of PGR5-deficient plants at constant daylight and their photosynthesis adapted to 380 µmol m^−2^ s^−1^ were largely unaffected, decreased PSI abundance suggested that edited plants experienced photodamage already in those conditions. Therefore, we conducted ROS staining to probe for oxidative stress in plants grown at constant daylight (Fig. S3). We utilised the general ROS marker H_2_DCFDA which fluoresces after reaction with H_2_O_2_ and other radicals such as OH· (Akter et al., 2021). Both Mes and BS cells of PGR5-deficients plants had higher levels of ROS, indicated by higher H_2_DCFDA fluorescence, already at growth light, compared to WT (Fig. 4f, g). After the exposure to high irradiance (30 min of 2000 µmol m^−2^ s^−1^), the ROS-responsive H_2_DCFDA signal intensity significantly increased in WT plants but not in plants lacking PGR5. No differences in a ratio of ROS levels between Mes and BS cells across treatments or genotypes were detected.

An oxidative stress in PGR5-deficient plants was also corroborated by results of RNAseq from leaves of plants grown at constant daylight. Analysis of transcript abundances revealed a high number of differentially expressed genes in PGR5-deficient plants relative to WT, including multiple genes involved in ROS metabolism and oxidative stress (Fig. S4). This was in sharp contrast to *S. viridis* plants lacking NDH, which showed a far more striking growth phenotype under the same growth conditions but had far fewer differentially expressed genes, compared to PGR5-deficient plants (Ermakova et al., 2024).

### Instantaneous responses of electron transport to fluctuating light

Given the strong effect of fluctuating daylight on growth and photosynthesis of PGR5-deficient plants, we next studied instantaneous responses to fluctuating light of plants grown under constant daylight. For this, a fluctuating light sequence of 400 μmol m^−2^ s^−1^ and 2000 μmol m^−2^ s^−1^ was applied to leaves during measurements. In WT plants, quantum yields of PSI (φ_I_, φ_ND_ and φ_NA_) were consistent throughout the measurements, with similar values between all high-light phases and between all growth-light phases (Fig. 5a-c). In contrast, φ_I_ and φ_NA_ in PGR5-deficient plants were progressively decreasing and φ_ND_ was progressively increasing with each subsequent high-light phase. By the end of the third growth-light phase φ_I_ was 36% lower in plants lacking PGR5, compared to WT, and by the end of the third high-light phase, plants lacking PGR5 had high φ_NA_ of 0.27 compared to 0 in the WT. Moreover, during the fluctuating light sequence, plants lacking PGR5 showed a progressive decline in the maximum P700^+^ signal (P_M_), reporting on the total amount of active PSI reaction centres, compared to WT, suggesting that, in the absence of PGR5, plants were receiving more photodamage at each high-light phase (Fig. 5d, e).

**Fig. 5.**
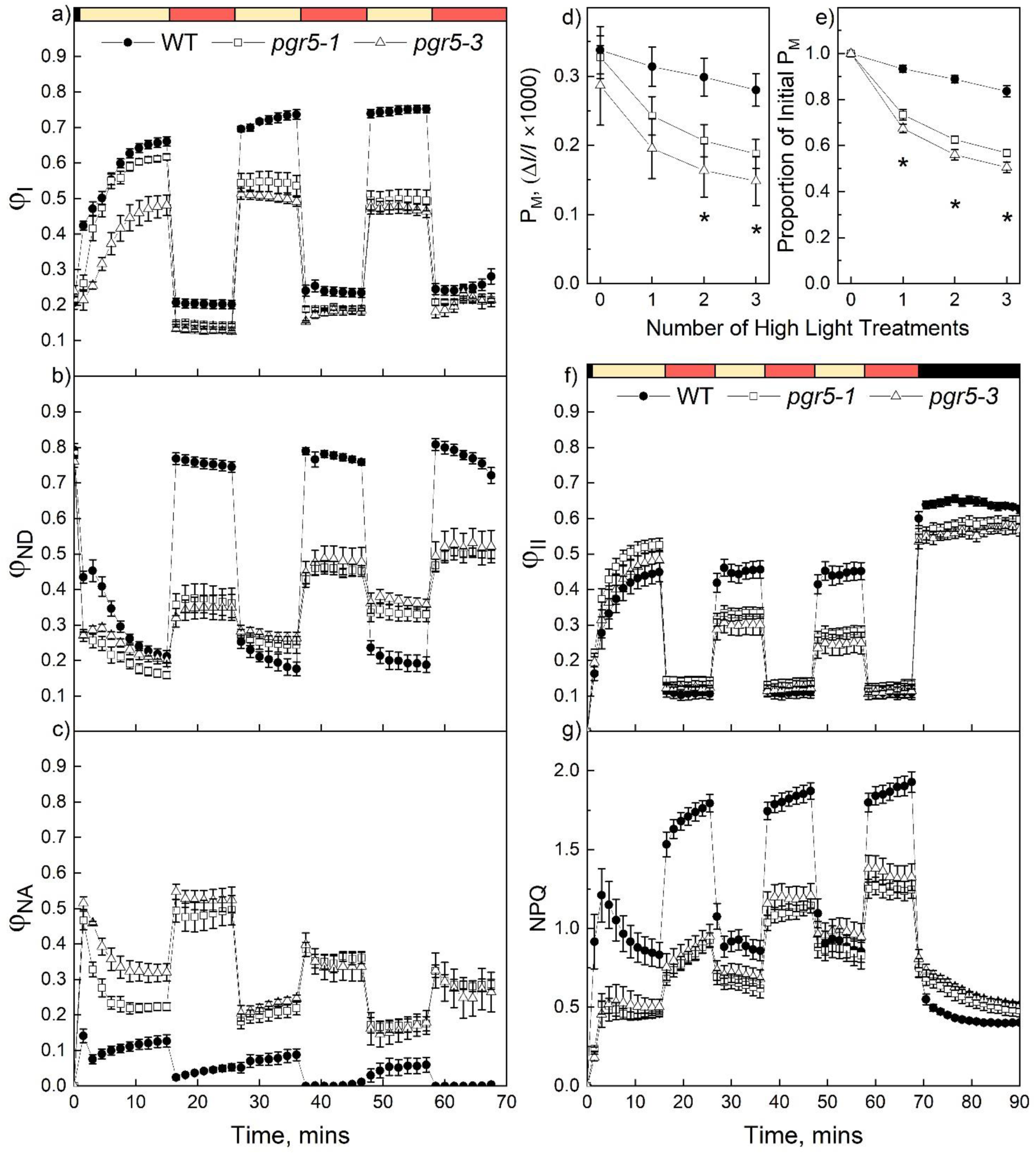
Instantaneous responses of Photosystem I (PSI) and Photosystem II (PSII) to fluctuating light in wild type (WT) *Setaria viridis* and plants lacking PGR5 (*pgr5-1* and *pgr5-3*) grown at constant daylight. (**a**) The quantum yield of PSI (φ_I_). (**b**) The non-photochemical yield of PSI due to donor side limitation (φ_ND_). (**c**) The non-photochemical yield of PSI due to acceptor side limitation (φ_NA_). (**d**, **e**) Changes in maximum photo-oxidisable P700 (PSI reaction centre), P_M_, monitored after each high-light treatment shown in (a - c). (**f**) The effective quantum yield of PSII (φ_II_). (**g**) Non-photochemical quenching (NPQ). Coloured bars indicate the irradiance during the measurement: black, 0 μmol m^−2^ s^−1^; yellow, 400 μmol m^−2^ s^−1^; red, 2000 μmol m^−2^ s^−1^. Mean ± SE, *n* = 4 biological replicates. Asterisks indicate significant differences between PGR5-deficient and WT plants (1-Way ANOVA with Tukey’s post hoc test at *P* < 0.05).

In regards to PSII activity, WT showed consistent φ_II_ values throughout the fluctuating light measurements which were similar between all high-light phases and between all growth-light phases (Fig. 5f). In contrast, φ_II_ in PGR5-deficient plants progressively decreased during each high-light period. WT and PGR5-deficient plants also showed some notable differences in NPQ when subjected to fluctuating light (Fig. 5g). WT plants built up NPQ to a similar level during each high-light phase and then relaxed it to a similar level during each growth-light phase. Contrastingly, plants lacking PGR5 were building up NPQ slower and reached the WT-levels only at the third growth-light stage. However, NPQ in PGR5-deficient plants was still significantly lower than in WT during the third high-light phase. Furthermore, relaxation of NPQ in darkness was drastically slower in the PGR5-deficient plants, compared WT, suggesting that PGR5-deficient plants engaged different NPQ mechanisms under fluctuating light (Fig. 5g).

Gas-exchange analysis of plants grown under constant daylight and subjected to fluctuating light sequence showed similar CO_2_ assimilation rates between each high-light phase and between each growth-light phase in the WT (Fig. S5). Comparatively, assimilation rates of PGR5-deficient plants were similar to WT during the first growth-light phase but progressively decreased during each subsequent change in irradiance.

### Membrane energisation and energy-dependent NPQ in PGR5-deficient plants

Next, we employed electrochromic shift (ECS) analysis to investigate whether inability of PGR5-deficient plants to promptly establish NPQ under sudden increases of irradiance were underpinned by changes in thylakoid membrane energisation. In C_4_ leaves, ECS probes proton fluxes specifically in Mes cells (Ermakova et al., 2024). Plants lacking PGR5 could only achieve about 60% of the WT *pmf* levels at all irradiances (Fig. 6a). Proton conductivity of the thylakoid membrane (*g*_H_^+^), reporting on ATP synthase activity, was significantly greater in PGR5-deficient plants than in the WT at 100 μmol m^−2^ s^−1^ and 2200 μmol m^−2^ s^−1^ (Fig. 6b). To estimate changes in CEF, *v*_H_^+^, the steady-state proton flux across the membrane, reflecting the combined activity of LEF and CEF was plotted against the relative electron flux through PSII, representing LEF alone, at corresponding irradiances (Fig. 6c) (Baker et al., 2007). In the absence of PGR5, plants showed a ∼21-25% decrease in slope, and thus in CEF.

**Fig. 6.**
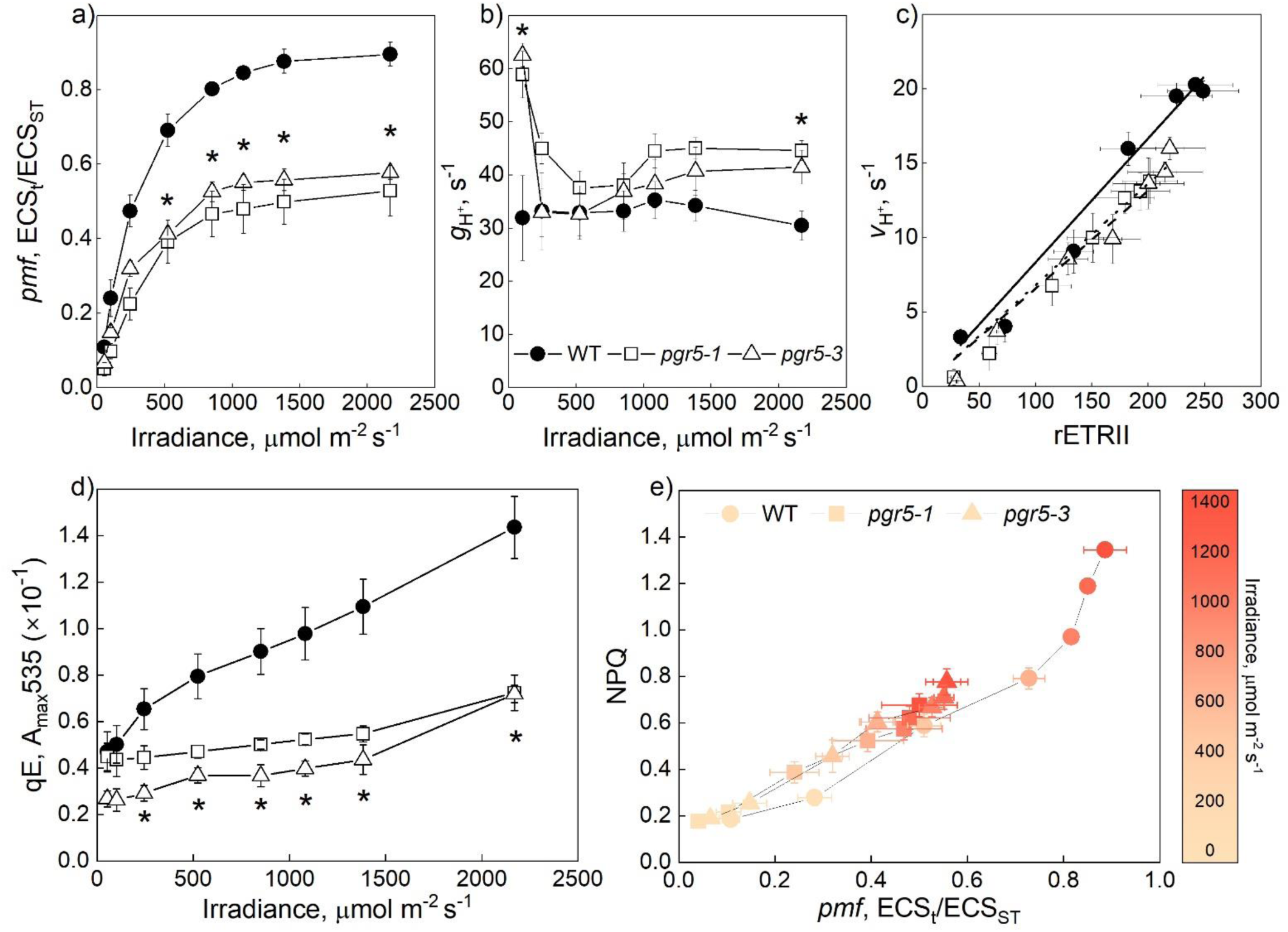
Thylakoid membrane energisation and non-photochemical quenching analysis in wild type (WT) *Setaria viridis* and plants lacking PGR5 (*pgr5-1* and *pgr5-3*) grown at constant daylight. (**a**, **b**) Proton motive force (*pmf*) and proton conductivity of the thylakoid membrane (*g*_H+_) after 3-min illumination at different irradiances. (**c**) The relationship between the light-driven proton flux (ν_H+_) and the relative Photosystem II electron transport rate (rETRII). (**d**) Leaf absorbance at 535 nm (qE) at different irradiances. (**e**) The relationship between non-photochemical quenching (NPQ) and *pmf* at corresponding irradiances. Mean ± SE, *n* = 3-5 biological replicates. Asterisks indicate significant differences between PGR5-deficient and WT plants (1-Way ANOVA with Tukey’s post hoc test at *P* < 0.05).

Since *pmf* has specific effects on the fast, energy-dependent NPQ, we probed leaf absorbance at 535 nm which is suggested to represent qE (Wilson et al., 2021). We compared the maximum leaf absorbance at 535 nm (A_max_535) reached at each light intensity between WT and PGR5-deficent plants (Fig. 6d). In the absence of PGR5, plants had drastically lower A_max_535 at irradiances above 100 μmol m^−2^ s^−1^ suggesting a strong deficiency in qE. Interestingly, when NPQ was plotted against *pmf* at corresponding irradiances (Fig. 6e), WT and PGR5-deficient plants showed a similar trajectory indicating that, overall, the NPQ response to *pmf* was unaltered in the absence of PGR5. However, plants lacking PGR5 could reach only about half of WT *pmf* and NPQ levels at irradiances above 500 μmol m^−2^ s^−1^ indicating that a lower NPQ in plants lacking PGR5 was a result of a limited capacity to generate *pmf*.

### Adaptations of PGR5-deficient plants to growth under fluctuating light

To further investigate the slowly relaxing NPQ observed in PGR5-deficient plants grown at constant daylight, we monitored NPQ build-up and relaxation in plants grown under fluctuating daylight during the fluctuating light cycles (Fig. 7a). WT plants showed distinct NPQ levels between growth-light and high-light which, however, were similar between all growth-light phases and between all high-light phases, indicating that NPQ was promptly responding to changes in irradiance. In contrast, plants lacking PGR5 showed lower amplitude of NPQ changes between growth-light and high-light phases and much slower responses to changes in irradiance but did show a gradual build-up of NPQ over time. Upon terminating the light, NPQ relaxed much slower in plants lacking PGR5 compared to WT.

**Fig. 7.**
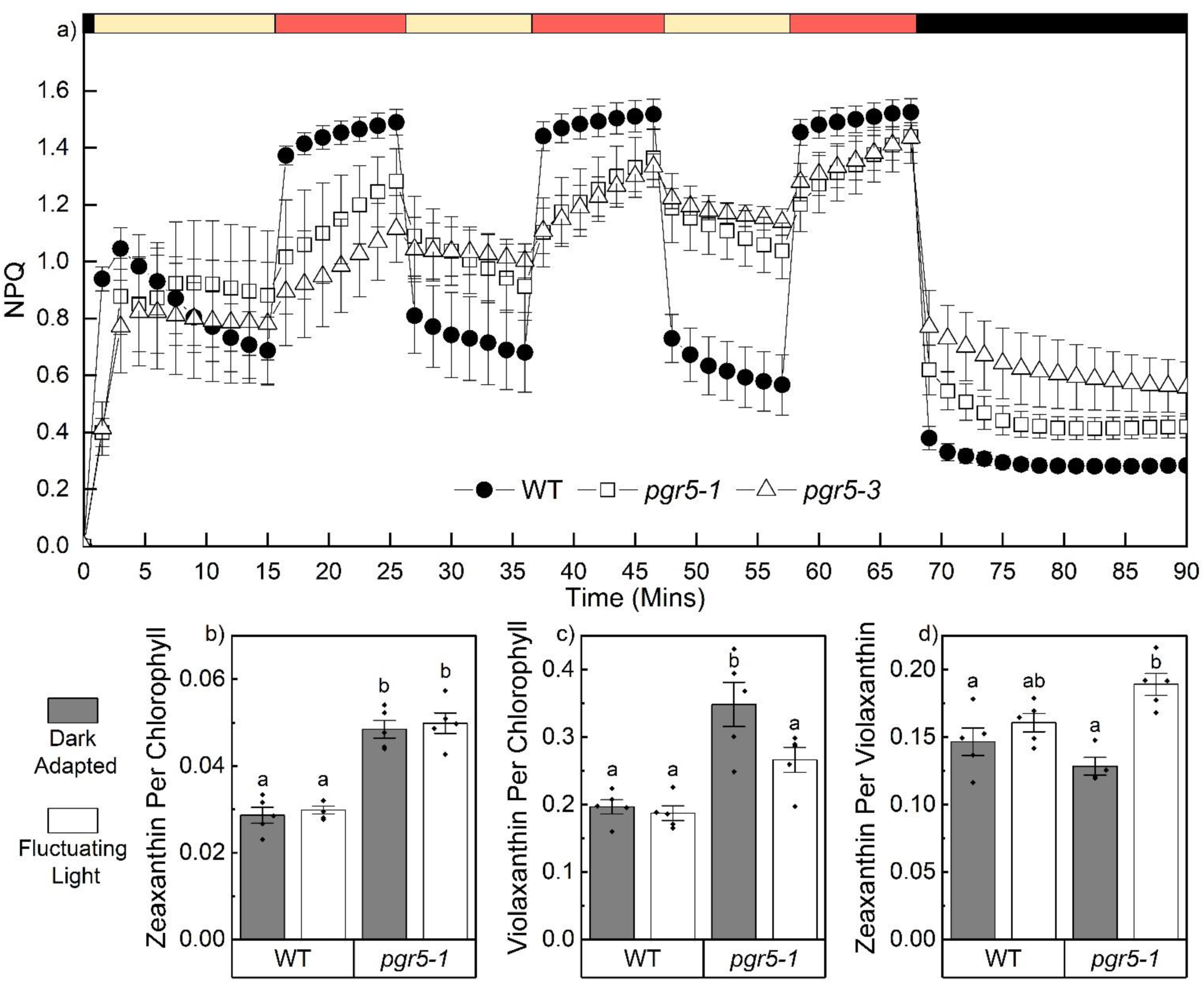
Analysis of non-photochemical quenching (NPQ) in wild type (WT) *Setaria viridis* and plants lacking PGR5 (*pgr5-1* and *pgr5-3*) grown at fluctuating daylight. (**a**) Dynamics of build-up and relaxation of NPQ during and after the fluctuating light sequence. Coloured bars indicate the irradiance during the measurement: black, 0 μmol m^−2^ s^−1^; yellow, 400 μmol m^−2^ s^−1^; red, 2000 μmol m^−2^ s^−1^. (**b** – **d**) Abundance of xanthophyll cycle pigments in leaves of dark- and fluctuating light-adapted plants. Mean ± SE, *n* = 4-5 biological replicates. Letters indicate significant differences between groups (2-Way ANOVA with Tukey’s post hoc test at *P* < 0.05).

To identify an origin of the slower-relaxing NPQ in PGR5-deficient plants, we analysed the content of xanthophyll pigments in leaves of plants grown under fluctuating daylight (Fig. 7b-d). Leaves were sampled following overnight dark adaptation and in the middle of the day during the low light phase of fluctuating light. Dark-adapted leaves of plants lacking PGR5 showed a significant increase in the relative abundance of zeaxanthin and violaxanthin per Chl, compared to WT. Under light, no significant changes in pigment abundance were detected in WT compared to the dark values. In contrast, a significant increase in the zeaxanthin/violaxanthin ratio was detected under light in plants lacking PGR5, compared to the dark values, indicating that, in the absence of PGR5 and qE, plants engaged the xanthophyll cycle to cope with fluctuating light.

## Discussion

In photosynthesis, the implementation of photoprotective mechanisms is essential for dissipating excess absorbed light energy which would otherwise result in the over-reduction of the electron transport chain, cause accumulation of ROS and result in photodamage (Murchie and Niyogi, 2011). In C_3_ species, PGR5 is a critical component of photoprotection and is indispensable when plants need to promptly upregulate qE in response to sudden increases of irradiance (Suorsa et al., 2012, Suorsa et al., 2016, Suorsa et al., 2013, Joliot and Johnson, 2011, Chaux et al., 2015, Yamamoto and Shikanai, 2019, Munekage et al., 2002). Abundance of PGR5 was elevated during the evolutionary transition from C_3_ to C_4_ photosynthesis and, in contrast to NDH, PGR5 is found in equal abundance in both Mes and BS cells of C_4_ plants (Ishikawa et al., 2016, Ermakova et al., 2021b, Munekage et al., 2010). Therefore, we have hypothesised that PGR5 is essential for photoprotection of C_4_ photosynthesis, likely in both cell types, and that an elevated abundance of PGR5 helped C_4_ plants inhabit high irradiance environments which are required to provide additional ATP for the C_4_ cycle (Sage et al., 1999, Hatch, 1987). To test this hypothesis, we generated gene-edited *S. viridis* lacking functional PGR5 (Fig. S1) and studied how the absence of PGR5 effected C_4_ photosynthesis and photoprotection.

A lack of PGR5 had more profound effects when plants were subject to changing irradiance compared to a constant light environment. Most prominently, plants lacking PGR5 grown at fluctuating daylight had greatly impaired growth and photosynthesis compared to constant daylight-grown plants (Fig. 1, 2, 3, 4). This severe fluctuating light growth phenotype was caused by a low CO_2_ assimilation capacity, which was not due to altered Rubisco abundance or CO_2_ delivery (Fig. 2b, 4d) but because of diminished abundances of photosynthetic complexes combined with the lower photochemical yields of photosystems (Fig. 3b, 4). Furthermore, plants lacking PGR5 grown at constant daylight showed an instant reduction of CO_2_ assimilation rate when exposed to high light (Fig. 2c). The lower NPQ at high irradiances in PGR5-deficient plants grown at constant daylight was likely a reason for this instantaneous high-light sensitivity of CO_2_ assimilation (Fig. 3c). As NPQ is critical for protecting photosynthetic machinery under light stress, PGR5 therefore plays a key role in photoprotection of C_4_ photosynthesis during changes of irradiance characteristic of plants’ natural habitats.

PGR5 is specifically important for the fast-relaxing form of NPQ, qE, which is regulated by lumen pH and therefore dependent on thylakoid membrane energisation. This is evident from the decreased NPQ in PGR5-deficient plants concomitant with the decreased *pmf,* as well as from a direct estimate of qE using absorbance at 535 nm (Fig. 6d, e). Moreover, in C_3_ plants PGR5 and qE are particularly important for safeguarding PSI under light stress (Munekage et al., 2008, Tiwari et al., 2016). Our results showed selective effects of the absence of PGR5 on PSI function also in a C_4_ plant. Both instantaneous and long-term damage to PSI was evident from the progressive decline of P_M_ with each next high-light phase of fluctuating light and from the drastically decreased abundances of PSI, Cyt*b*_6_*f* and NDH in PGR5-deficient plants grown under fluctuating daylight, while PSII abundance was less impacted (Fig. 4d, 5e). As NDH is predominantly found in BS cells and likely forms a supercomplex with PSI for stability, the lower abundance of NDH in PGR5-deficient plants could be attributed to a loss of PSI to BS cells (Ermakova et al., 2024, Kato et al., 2018). However, the patterns of ROS accumulation suggested that PSI damage occurred in both Mes and BS cells, which is consistent with the equal distribution of PGR5 in both cell types (Fig. 4h) (Ermakova et al., 2024, Ermakova et al., 2021b). Therefore, PGR5 is needed for PSI photoprotection in both types of cells in C_4_ leaves. Given the low content of PSII in BS cells, this raises many questions about how photoprotection operates in BS cells, which could be addressed in future studies.

Curiously, whilst having diminished qE capacity, plants lacking PGR5 showed increased NPQ when grown under fluctuating daylight (Fig. 1f, 3f). Though we cannot exclude that elevated abundance of PsbS detected in the PGR5-deficient plants in these conditions (Fig. 4d) could contribute to upregulating qE, our results suggest that PGR5-deficient plants upregulated slow-relaxing NPQ mechanisms to cope with light stress. Both constant daylight- and fluctuating daylight-grown PGR5-deficient plants showed the slow build-up of NPQ over multiple high-light phases as well as slow NPQ relaxation (Fig. 5g, 7a). In addition, leaves of plants lacking PGR5 contained more zeaxanthin and violaxanthin when grown in fluctuating daylight and showed increased conversion of violaxanthin to zeaxanthin during the day (Fig. 7b-d). The timescale of these NPQ responses and the increased capacity and activity of the xanthophyll cycle are indicative of a slower interconverting pool of xanthophylls involved in qZ being engaged in PGR5-deficient plants (Jahns and Holzwarth, 2012, Nilkens et al., 2010). In the absence of qE, qZ becomes the next best option for NPQ. However, due to the slow nature of qZ, under fluctuating light, the constitutively high NPQ leads to significant productivity losses during low-light phases. This high-quenching state further explains the severe growth and photosynthesis phenotype of the PGR5-deficient plants under fluctuating daylight (Fig. 1). Moreover, as long-term high-light responses do not rely on qE (Ruban, 2016, Jahns and Holzwarth, 2012, Nilkens et al., 2010), growth and photosynthesis of plants lacking PGR5 was not affected under a high constant daylight (Fig. S3). These results highlight the essential function of qE and PGR5 in maintaining high assimilation rates of C_4_ photosynthesis under fluctuating light.

While the role of PGR5 in building up *pmf* to regulate qE is well-established in C_3_ plants and confirmed in a C_4_ plant in this work, the exact mechanism of PGR5’s contribution to *pmf* is still uncertain. It is proposed that PGR5 builds up *pmf* either through regulating CEF around PSI or through minimising *pmf* dissipation by lowering proton conductivity of ATP synthase (Kanazawa et al., 2017, Yamori and Shikanai, 2016). Although the proton conductivity (*g*_H_^+^) was slightly increased in the absence of PGR5, our results point more to a function of PGR5 in regulating CEF. Strong PSI acceptor side limitation in plants lacking PGR5 (Fig. 3i, l), was consistent with the lack of CEF accepting electrons from PSI. Furthermore, estimating CEF by plotting the light-driven proton flux across the thylakoid membrane (*v*_H_^+^) against rETRII (Avenson et al., 2005) also showed suggested lower CEF in the absence of PGR5 (Fig. 6c). It is worth mentioning here that both ECS and fluorescence analysis of *S. viridis* leaves predominantly report on Mes cells due to BS cells having low PSII abundance and activity and lacking ECS signal (Ermakova et al., 2024). The 21-25% decrease in slope observed in PGR5-deficient *S.viridis* is consistent with the analyses of the Arabidopsis *pgr5* mutant, suggesting a similar mechanism for PGR5 function in C_3_ and C_4_ Mes cells (Avenson et al., 2005, Livingston et al., 2010, Degen et al., 2023).

Overall, our results indicate that the role of PGR5 in photoprotection of C_4_ photosynthesis is consistent with the function of PGR5 in C_3_ plants (Suorsa et al., 2012, Suorsa et al., 2016, Joliot and Johnson, 2011, Yamamoto and Shikanai, 2019, Munekage et al., 2002, Avenson et al., 2005). Although it is hard to make direct comparison between C_3_ and C_4_ species since the severity of fluctuating light can vary for each species depending on exact cycle parameters, it does seem likely that growth of C_3_ *pgr5* mutants under fluctuating daylight is more severely affected, compared to *S. viridis* lacking PGR5, as fluctuating light is often lethal for C_3_ *pgr5* mutants (Tikkanen et al., 2010, Suorsa et al., 2012, Long et al., 2022, Niu et al., 2023, Munekage et al., 2002). This suggests that the importance of PGR5 and qE may differ between C_3_ and C_4_ photosynthesis, possibly due to the function of PGR5 being partially complemented through upregulation of NDH in BS chloroplasts (Livingston et al., 2010). In addition, although PSII in Mes cells produces most of the reducing power required for the Calvin cycle, C_4_ Mes electron transport is not directly regulated by the Calvin cycle activity like in C_3_ Mes cells. While the C_4_ metabolic cycle is suggested to make C_4_ photosynthesis slower in reacting to changes in environment (Kubasek et al., 2013, Li et al., 2021), the cycle, and also the electron transport chain of BS cells, may serve as a buffer to ‘absorb’ any imbalances between Mes electron transport supply and Calvin cycle demand, potentially presenting an additional advantage of the two-cell C_4_ system.

## Conclusion

Through the analysis of gene-edited *S. viridis* with new null *pgr5* alleles, our work demonstrates that PGR5 is essential for generating *pmf* and regulating qE to facilitate photoprotection in C_4_ plants. Even at moderate growth irradiance PGR5-deficients plants experience oxidative stress, while growth under fluctuating light results in severe photodamage, impairing growth and photosynthesis. However, in the absence of PGR5 and qE, *S. viridis* upregulates qZ as an alternative photoprotective mechanism. This work uncovers the role of various NPQ mechanisms in C_4_ plants and thus provides novel insights into dynamic regulation of C_4_ photosynthesis in naturally fluctuating light environments and informs potential strategies for improving C_4_ crops.

## Materials and Methods

### Generation of *S. viridis* lacking PGR5

*S. viridis* cv. ME034V-1 plants with null *pgr5* alleles were created using CRISPR/Cas9 gene-editing as described in Ermakova et al. (2024). The genomic and coding sequences of *S. viridis pgr5* (Sevir.6G259600) were obtained from Phytozome (https://phytozome-next.jgi.doe.gov). Cas9 was targeted to the first exon of *pgr5* using gRNAs ACGTGGTCCAGCTCCGTGTC and AGACCGGGGCCGCGCCGAC (Fig. S1). gRNAs were selected using CRISPOR (Concordet and Haeussler, 2018) and assembled into a synthetic polycistronic gene, according to Xie et al. (2015). The Golden Gate cloning system (Engler et al., 2014) was used to assemble the construct which was transformed into *S. viridis* using the *Agrobacterium tumefaciens* strain AGL1 (Osborn et al., 2017). Editing of the target gene was confirmed by sequencing in T_1_ plants and the progenies of plants with homozygous null alleles were used in all experiments. Four distinct null *pgr5* alleles were obtained and designated *pgr5-1*, *pgr5-2*, *pgr5-3* and *pgr5-4* (Fig. S1). Gene and protein sequences were visualised in Geneious Prime (Dotmatics, https://www.geneious.com/). Of the four null alleles, *pgr5-1* and *pgr5-2* resulted in a similar amino acid sequence as did *pgr5-3* and *pgr5-4*. Therefore, *pgr5-1* and *pgr5-3* were mainly used in experiments.

### Plant growth conditions

*S. viridis* plants were germinated either on rooting medium described in Osborn et al. (2017) or directly in 0.5 L pots with seed raising mix (Debco, Tyabb, Australia) containing 5 g L^−1^ of Osmocote fertiliser (Scotts, Bella Vista, Australia). When germinated on rooting medium, plants were transferred to soil in 0.5 L pots one week after germination. When germinated on soil, seeds were covered with inverted plastic cups to maintain humidity, with the cups removed one week after germination. In both cases plants were grown in a controlled-environment chamber with a 16 h light/8 h dark photoperiod, an irradiance of 380 µmol m^−2^ s^−1^, ambient CO_2_, 28°C day, 22°C night and 60% humidity (referred to as constant daylight). The position of WT and edited plants within the chamber was blocked and randomised to minimise positional effects. Plants were analysed 3-4 weeks after sowing, with the youngest fully expanded leaves used for all analyses. WT *S. viridis* plants were used as a control for all experiments.

For fluctuating light growth experiments, plants were germinated on soil under constant daylight as described above. One week post-germination, plants were moved to a chamber with 5 minute cycles of 250 µmol m^−2^ s^−1^ followed by 1 minute at 1000 µmol m^−2^ s^−1^, repeating for an entire 16 hr photoperiod and giving a total integrated irradiance of 375 µmol m^−2^ s^−1^ corresponding to constant daylight conditions. For high constant daylight experiment, plants were grown at an irradiance of 1000 µmol m^−2^ s^−1^ over a 16 h light/8 h dark photoperiod.

### Chlorophyll quantification and immunoblotting

For analysis of Chl content and photosynthetic proteins, 0.79 cm^2^ leaf discs were flash-frozen in liquid nitrogen. Chl was extracted in 80% acetone buffered with 25 mM HEPES (pH 7.8), and Chl *a* and *b* content was determined according to Porra et al. (1989).

Protein extraction from leaf discs, SDS-PAGE and immunoblotting was performed as described in Ermakova et al. (2019). Samples were normalised on leaf area basis, separated by SDS-PAGE and transferred to nitrocellulose membrane. Samples were probed with antibodies against key photosynthetic proteins: PGR5 (1:3000, AS163985, Agrisera, Vännäs, Sweden), NdhH (NDH, 1:3000, AS164065, Agrisera), Rieske (Cyt*b*_6_*f*, 1:5000, AS08330, Agrisera), RbcL (Rubisco, Martin-Avila et al. (2020)), PsaB (PSI, 1:5000, AS10 695, Agrisera), D1 (PSII, 1:10,000, AS10704, Agrisera), AtpB (ATP synthase, 1:10,000, Agrisera), PsbS (1:3000, AS09533, Agrisera). Immunoblots were imaged and quantified using ChemiDoc MP Imaging System and Image Lab software (Bio-Rad, Hercules, CA, USA).

### ROS staining, microscopy and quantification

The accumulation of ROS in *Setaria* leaves was probed using 2′,7′-dichlorofluorescein diacetate (H_2_DCFDA) which fluoresces following reaction with reactive oxygen species such as hydrogen peroxide (H_2_O_2_) and hydroxyl radicals when excited between 460 – 500 nm. Using methods adapted from Fichman et al. (2019) and Zhu et al. (2016), ∼8 cm long leaf segments were harvested from plants adapted to growth irradiance (380 µmol m^−2^ s^−1^) or following a 30-min high irradiance (2000 µmol m^−2^ s^−1^) treatment. Following sample collection, leaves were vacuum infiltrated with 500 μM H_2_DCFDA in 50 mM sodium phosphate buffer (pH 7.4) for 5 minutes. Immediately following infiltration, leaves were hand-sectioned into ∼0.5 mm wide cross-sections using a razor blade. Leaf sections were then placed onto a glass slide, mounted in water and imaged under a Leica DM5500 fluorescence microscope using the brightfield channel to capture leaf anatomical features, the 527/30 nm bandpass filter to capture H_2_DCFDA fluorescence and the 700/75 nm filter to capture Chl *b* fluorescence.

Following imaging, ROS abundance and localisation were quantified using LasX (Leica Microsystems). For each leaf section, the H_2_DCFDA fluorescence signal was quantified for individual wreath units each consisting of the vascular bundle surrounded by BS and Mes cells. Quantification of total ROS per wreath was determined by selecting an entire wreath as a region of interest (ROI). The fluorescence signal of BS cells was then quantified by selecting the central part of each wreath encompassing the BS and vasculature as a second ROI. The Mes signal was then calculated by subtracting the fluorescence signal in the BS ROI from the total fluorescence signal of the whole wreath ROI.

### Leaf spectroscopic and fluorescence analyses

Simultaneous measurements of Chl fluorescence and the redox state of P700 (the reaction centre of PSI) were performed using DUAL-PAM/F (Heinz Walz, Effeltrich, Germany) under a red actinic light (635 nm) using saturating pulses of red light (300 ms, 12,000 μmol m^−2^ s^−1^). The redox state of P700 was assessed by detecting absorbance changes of the P700^+^ cation at 830 nm with a dual wavelength unit (830/875 nm). Leaves were dark-adapted for 30 minutes before a saturating pulse was applied to obtain the maximum (F_M_) and minimum (F_0_) levels of fluorescence. F_V_/F_M_, the maximum quantum yield of PSII, was calculated as (F_M_ - F_0_) / F_M_. Following this, the maximal P700^+^ signal (P_M_) was recorded upon the application of a saturating pulse at the end of an 8-second far-red light (720 nm) illumination, with the minimal P700^+^ signal (P_0_) recorded directly after the saturating pulse. Following the determination of dark parameters leaves were light acclimated to 480 µmol m^−2^ s^−1^ for 15 minutes.

Next, light response curves were measured by progressively increasing the actinic light from 55 to 1650 µmol m^−2^ s^−1^ and applying a saturating pulse at the end of every 120-s illumination period. Upon the application of each saturating pulse, F_M_’ (the maximum level of fluorescence under light) and F (the fluorescence level before the application of a saturating pulse) were recorded. This allowed for the tracking of the partitioning of absorbed light energy within PSII between the photochemical (φ_II_ = (F_M_’ – F) / F_M_’) and non-photochemical reactions, including the regulated (φ_NPQ_ = (F_M_ – F ^’^) / F_M_) and non-regulated (φ_NO_ = F / F_M_) fractions (Kramer et al., 2004). NPQ was calculated after Bilger and Björkman (1990) as (F_M_ – F_M_’) / F_M_’ and represents the combined contributions of qE, qZ and qI. The photochemical yield of PSI (φ_I_ = (P_M_’ – P) / (P_M_ – P_0_)), the non-photochemical yields of PSI due to acceptor (φ_NA_ = (P_M_ – P_M_’) / (P_M_ – P_0_)) or donor (φ_ND_ = (P – P_0_) / (P_M_ – P_0_)) side limitation were calculated as described in Klughammer and Schreiber (2008).

### Electron transport analyses under fluctuating light

Electron transport analyses under fluctuating light were performed by changing the incident irradiance on the leaf between darkness (0 µmol m^−2^ s^−1^), ambient irradiance (400 µmol m^−2^ s^−1^) and high irradiance (2000 µmol m^−2^ s^−1^) over the course of the measurement. For Chl fluorescence analysis, plants were assayed in a LICOR-6800 IRGA (LI-COR Biosciences) equipped with a 6800-01A fluorometer head. For all measurements 90% red/10% blue actinic light was used with 25°C leaf temperature, 55% relative chamber humidity, flow rate of 500 µmol s^−1^, flow pressure of 0.1 kPa, fan speed of 10,000 RPM and CO_2_ partial pressure of 400 μmol mol^−1^ on the reference side. At the beginning of each measurement, a saturating fluorescence pulse was applied to a dark-adapted leaf to obtain F_0_ and F_M_. Saturating pulses were then applied every 90 seconds over the course of the fluctuating actinic light sequence and in darkness. NPQ was calculated upon the application of each saturating pulse as described earlier.

Measurements of P700 redox state were performed using a DUAL-PAM/F under a red actinic light (635 nm). Following dark adaptation, the P_M_ and P_0_ were recorded as described earlier. Then, saturating pulses were applied every 90 seconds over the course of the fluctuating actinic light sequence. PSI electron transport parameters were calculated upon the application of each saturating pulse as described earlier. Additionally, to analyse the extent of PSI damage resulting from exposure to high irradiance, the P_M_ and P_0_ were recorded again at the end of each high-light period.

### Gas-exchange analysis

All gas exchange analyses were performed using the LICOR-6800 equipped with a 6800-01A fluorometer head. For CO_2_ response curves, leaves were clamped into the chamber and equilibrated at 400 μmol mol^−1^ CO_2_ on the reference side and 500 μmol photons m^−2^ s^−1^ for 15 minutes. CO_2_ assimilation rates were then recorded at CO_2_ partial pressures from 0 to 1600 μmol mol^−1^ at 3-minute intervals. For light response curves, leaves were equilibrated at 400 μmol mol^−1^ CO_2_ and 500 μmol m^−2^ s^−1^ irradiance for 15 minutes and then CO_2_ assimilation was recorded first at irradiance progressively increasing from 0 to 3000 μmol m^−2^ s^−1^ and then decreasing from 3000 to 0 μmol m^−2^ s^−1^. Measurements were made at 3-minute intervals. CO_2_ and light response curves were fit using the biochemical model of C_4_ photosynthesis parameterised for *S. viridis* (von Caemmerer, 2021). The quantum yield of CO_2_ assimilation (Φ), light compensation point (LC) and dark respiration rate (*R*_d_) were determined by fitting a linear regression through the initial slope of light response curves (from 0 to 100 μmol m^−2^ s^−1^) in Excel 2016 (Microsoft).

### Thylakoid Membrane Energisation

Electrochromic shift (ECS) signal and absorbance at 535 nm were monitored with the Dual PAM-100 equipped with the P515/535 emitter-detector module. ECS was estimated from the absorbance changes between 550 and 520 nm and normalised for the amplitude of ECS response to a saturating pulse (20 μs, 14 000 μmol m^−2^ s^−1^) measured from the dark-adapted leaves (ECS_ST_). Changes in ECS signal were measured during 3-minute illumination periods of increasing irradiance. After each illumination period, the light was switched off for 20 seconds and the kinetics of ECS decay were analysed to determine *pmf*, proton conductivity of the thylakoid membrane (*g*_H_^+^) and the light-driven proton flux across the thylakoid membrane (*v*_H_^+^). *Pmf* was estimated as the total change in amplitude of ECS signal upon light-to-dark transition (ECS_t_) normalised for ECS_ST_ (Takizawa et al., 2007). *g*_H_^+^ was calculated as the inverse time constant of the first-order exponential decay in ECS signal upon the termination of light (Sacksteder et al., 2000) and was fit in OriginPro 2023b (OriginLab, Northampton, MA, USA). *v*_H_^+^ was calculated as the instantaneous initial rate of change in ECS signal upon light termination, also fit in OriginPro 2023b (Takizawa et al., 2008).

The absorbance signal at 535 nm is informative of lumen acidification, zeaxanthin conversion and qE (Schreiber and Klughammer, 2008). The absorbance at 535 nm was measured in a separate experiment, during 3-minute illumination periods of increasing irradiance. The highest 535 nm absorbance value reached at each illumination period was recorded as the maximum absorbance (A_max-_535) for that irradiance.

### Xanthophyll analysis

To obtain samples for xanthophyll analysis, leaf discs were taken from plants grown under fluctuating daylight conditions. Leaves were sampled both following overnight dark adaptation and in the middle of the day during the low-light phase (250 µmol m^−2^ s^−1^) of the fluctuating daylight sequence. After sampling, leaf discs were flash-frozen in liquid nitrogen and then freeze-dried. Pigments were extracted using acetone-ethyl acetate (6:4, v/v) as previously described in Alagoz et al. (2020). Prior to separation, 100 µg of α-tocopheryl acetate (Sigma) was added to each sample as an internal standard. Pigments were then separated on an Agilent 1200 series high-performance liquid chromatograph (HPLC) (Agilent Technologies) using a C30 (4.6 x 250 mm; YMC) column coupled to a 5 µm (4.0 x 23 mm) C30 guard column. 40 µl sample injections were separated at a flow rate of 1.0 mL min^−1^ with mobile phases of 100% methanol and 100% methyl tert-butyl ether, with a gradient elution performed as previously described in Alba et al. (2005).

### Bulk RNA-seq analysis

Raw mRNA-sequencing reads were mapped to the *S. viridis* genome utilizing the Star Seq Toolkit (Dobin et al., 2013). The raw reads were assessed and trimmed using fastqc version 0.11.9 and trim galore 0.6.7 with low-quality base calls below a PHRED score of 20 trimmed by applying the parameter -q 20. Transcript quantification was conducted for unique matches at gene level using StringTie against the *S. viridis* v 4.1. The resulting raw counts were imported into edgeR, focusing on loci detected at counts per million (CPM) exceeding 1 in a minimum of three samples. Library normalisation was performed using voom, employing the trimmed mean of M-values (TMM) method to account for both sequencing depth and composition.

### Statistical Analysis

1-Way and 2-Way ANOVAs were performed in OriginPro 2023b using a significance threshold of *P* < 0.05. Details of replication, post hoc tests and *P* values are provided in the main text and in figure legends.

## Acknowledgements

We thank Prof Spencer Whitney for a gift of vectors and antibodies, Dr Xueqin Wang for Setaria transformation, and Emily Watson for technical assistance. We thank the Australian Plant Phenomics Network supported under the National Collaborative Research Infrastructure Strategy of the Australian Government. This work was supported by the Australian Research Council (CE140100015).

## Author contributions

ME, RTF and RW designed the research; ME, RTF, SC, KXC and BP supervised research; ME generated gene-edited plants; RW, JW, MM, SY and SJN performed experiments; RW, ME, MM, JW, RTF and SC analysed data; RW and ME wrote the paper; all authors discussed the results and contributed to the final manuscript.

