## Supplementary Materials for "PGR5 promotes energy-dependent non-photochemical quenching to enable efficient C_4_ photosynthesis under fluctuating light"

**Table S1**. Photosynthetic parameters of WT *S. viridis* and PGR5-deficient plants (*pgr5-1* and *pgr5-3*) fit from the light curves of $CO_2$ assimilation (Fig. 2c). $R_d$, respiration in the dark; LC, light compensation point; Φ, quantum yield of $CO_2$ assimilation (see details of fitting in the Materials and Methods section). Asterisks indicate statistically significant differences between the PGR5-deficient plants and WT (1-Way ANOVA with Tukey's post hoc test at $P < 0.05$). Mean ± SE, $n$ = 4-5 biological replicates.

| **Parameter** | Constant Daylight | | | Fluctuating Daylight | | |
|---|---|---|---|---|---|---|
| | **WT** | ***pgr5-1*** | ***pgr5-3*** | **WT** | ***pgr5-1*** | ***pgr5-3*** |
| $R_d$ (μmol $CO_2$ $m^{-2}$ $s^{-1}$) | 1.87 ± 0.31 | 1.67 ± 0.19 | 1.88 ± 0.29 | 1.82 ± 0.25 | 0.83 ± 0.12* | 0.55 ± 0.24* |
| LC (μmol irradiance $m^{-2}$ $s^{-1}$) | 27.96 ± 2.58 | 30.42 ± 1.62 | 33.82 ± 3.05 | 26.94 ± 1.22 | 51.30 ± 7.73* | 46.46 ± 12.55 |
| Φ (Increasing Irradiance) (mol $mol^{-1}$) | 0.0663 ± 0.005 | 0.0551 ± 0.004 | 0.0550 ± 0.004 | 0.0674 ± 0.007 | 0.0148 ± 0.002* | 0.0078 ± 0.004* |
| Φ* (Decreasing Irradiance) (mol $mol^{-1}$) | 0.0569 ± 0.004 | 0.0301 ± 0.004* | 0.0292 ± 0.002* | 0.0569 ± 0.002 | 0.0076 ± 0.003* | 0.0069 ± 0.003* |

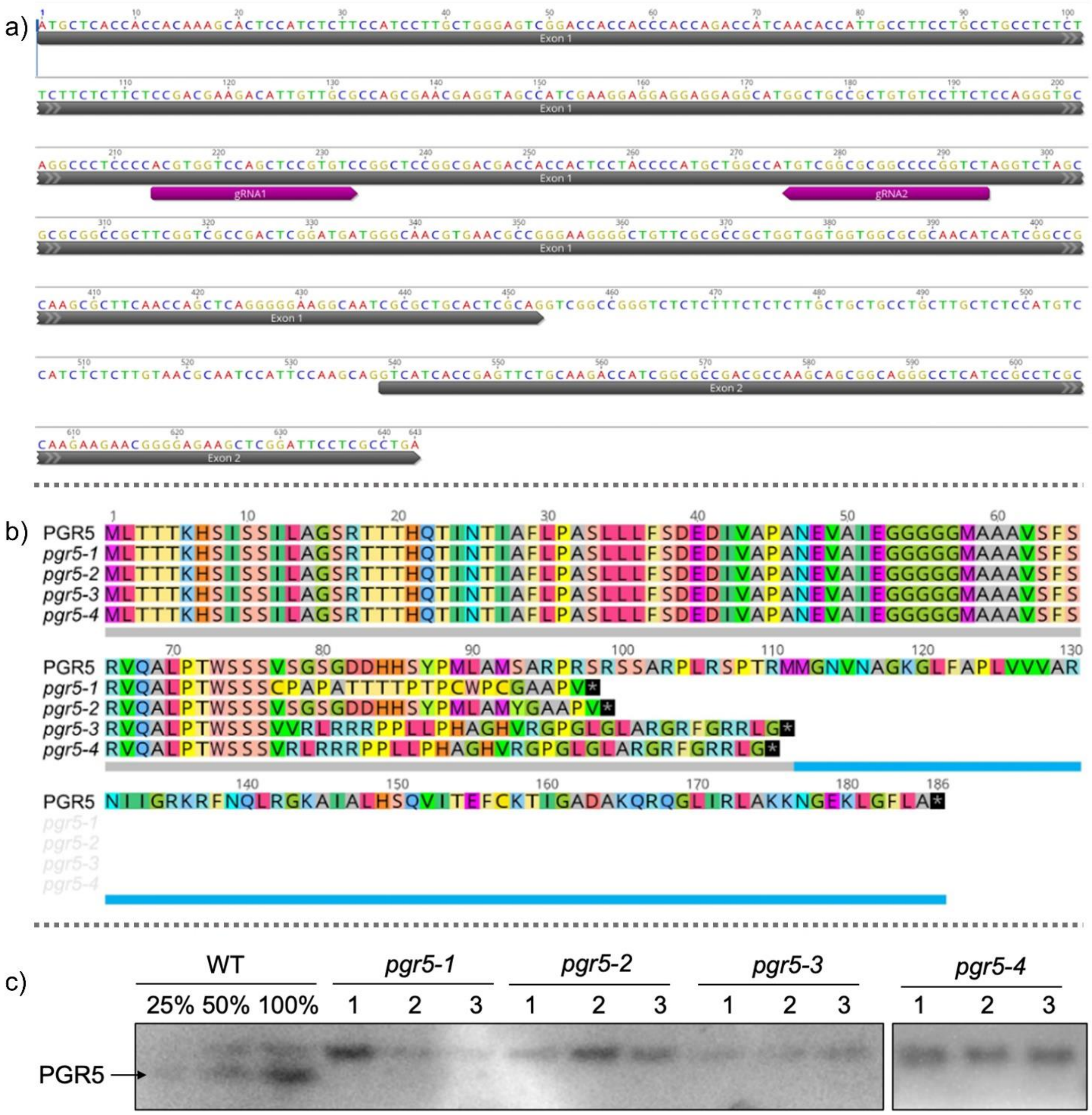


**Fig. S1.** Four new *pgr5* alleles obtained via CRISPR/Cas9 gene-editing of *S. viridis*. (**a**) Location of guide RNAs (gRNAs) targeting the first exon of the *pgr5* genomic sequence. (**b**) Amino acid sequences of PGR5 and proteins predicted to be encoded by the new *pgr5* alleles*.* Grey bar underneath indicates chloroplast signal peptide, blue bar – mature protein. (**c**) Immunoblotting of leaf protein extracts from WT and plants homozygous for the edited alleles (*pgr5-1*, *pgr5-2*, *pgr5-3* and *pgr5-4)* with PGR5-specific antibodies. The *pgr5-4* blot is modified from (Ermakova et al., 2024).

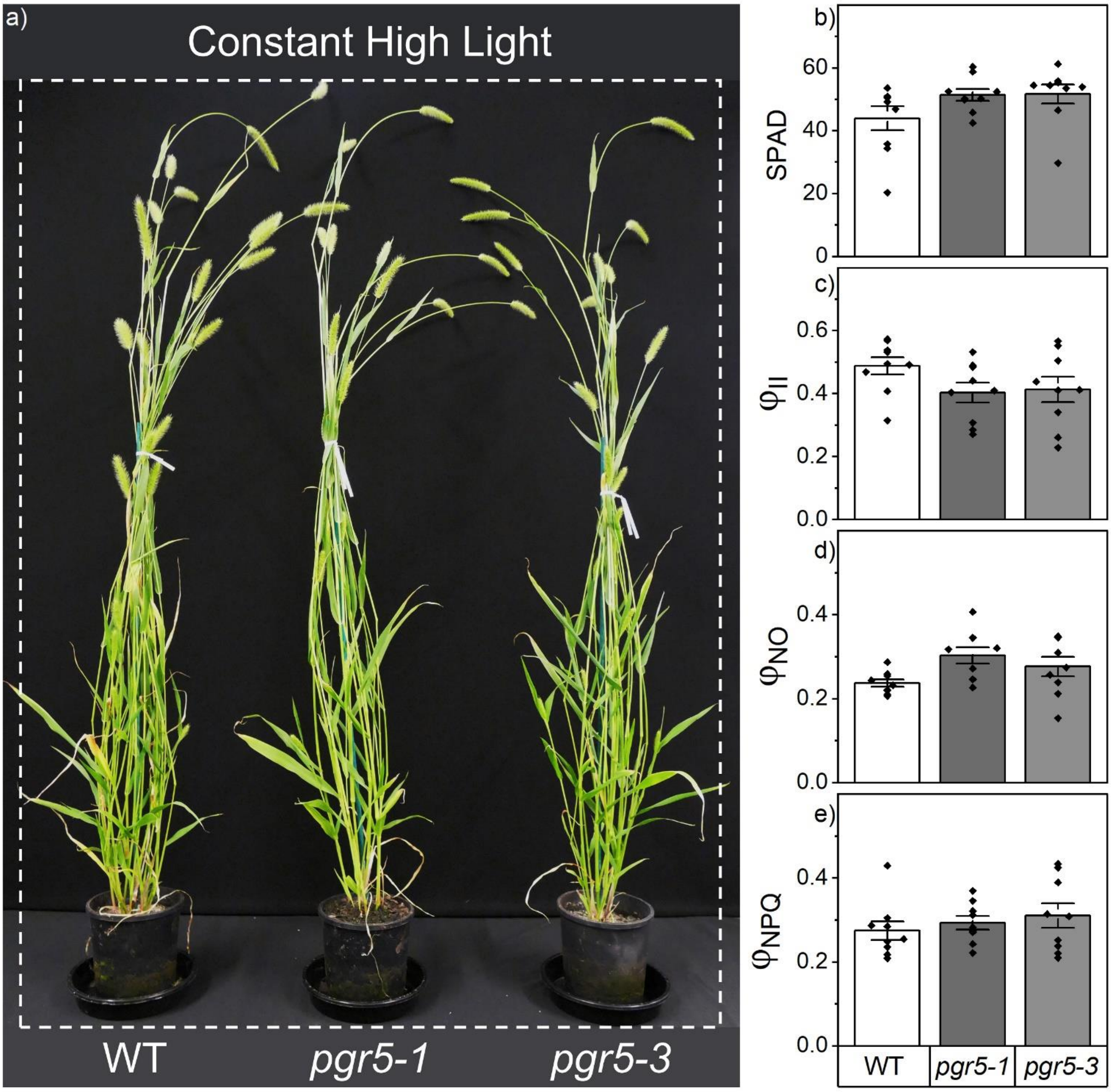


**Fig. S2**. Growth and photosynthesis of WT and PGR5-deficient plants grown under high constant daylight (1000 µmol $m^{-2}$ $s^{-1}$). (**a**) Growth phenotype of plants 45 days after germination. (**c-e**) Leaf photosynthesis parameters analysed with MultispeQ: relative chlorophyll content (SPAD), the effective quantum yield of PSII ($\varphi_{II}$), the yield of non-regulated non-photochemical reactions ($\varphi_{NO}$) and the yield of non-photochemical quenching ($\varphi_{NPQ}$). Mean ± SE, *n* = 9 biological replicates. No significant differences were found between WT plants and plants with null *pgr5* alleles. (1-Way ANOVA with Tukey's post hoc test at $P < 0.05$).

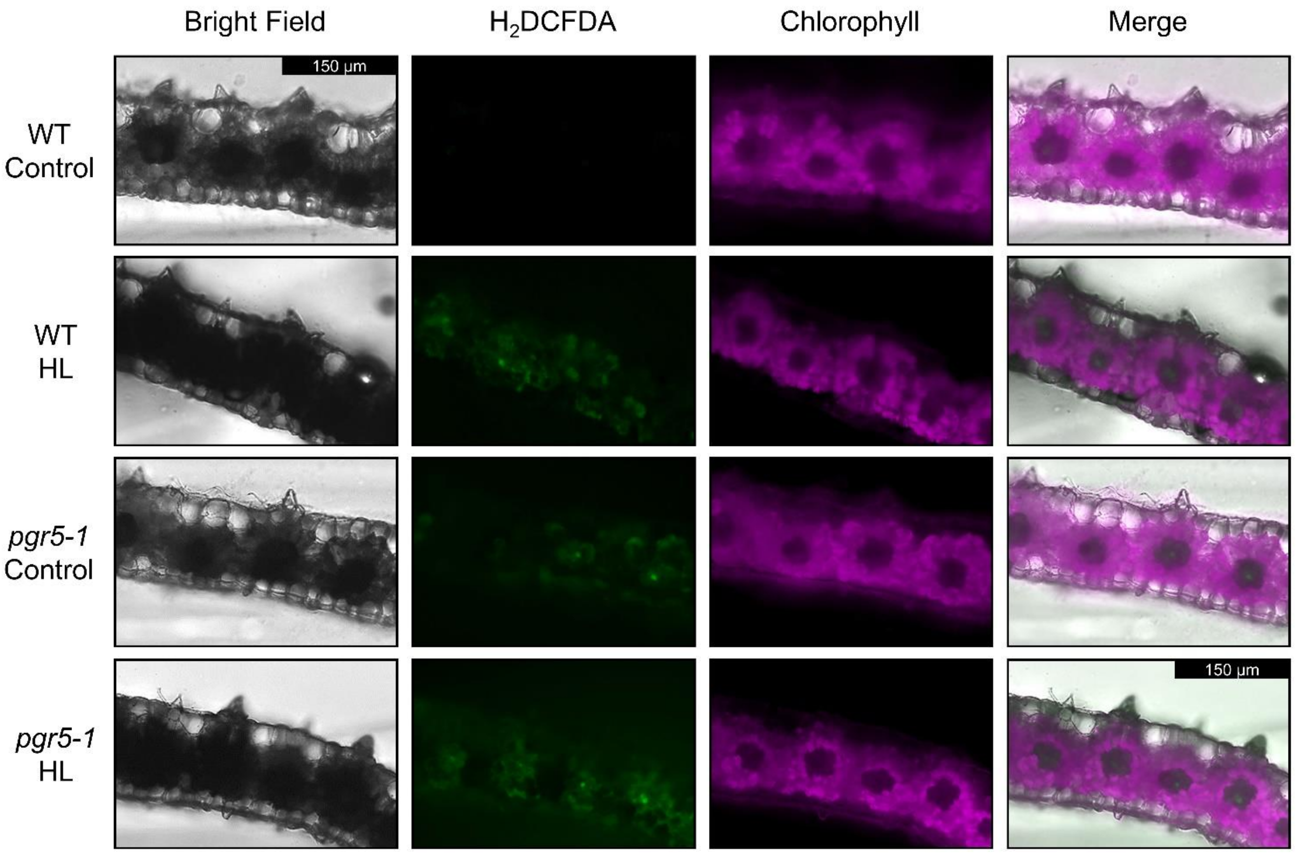


**Fig. S3**. Representative microscopy of ROS staining in WT and *pgr5 Setaria viridis* under growth light (control) and after a 30-minute high-light treatment (HL). Bright field images show the structure of leaf sections. $H_2$DCFDA fluorescence (green) shows the accumulation of ROS following reaction of the dye with $H_2O_2$. Chlorophyll fluorescence (magenta) demonstrates the localisation of chlorophyll associated with PSII, and therefore Mes cells in the section. A merge of the three channels demonstrates the localisation of ROS and PSII within the leaf section.

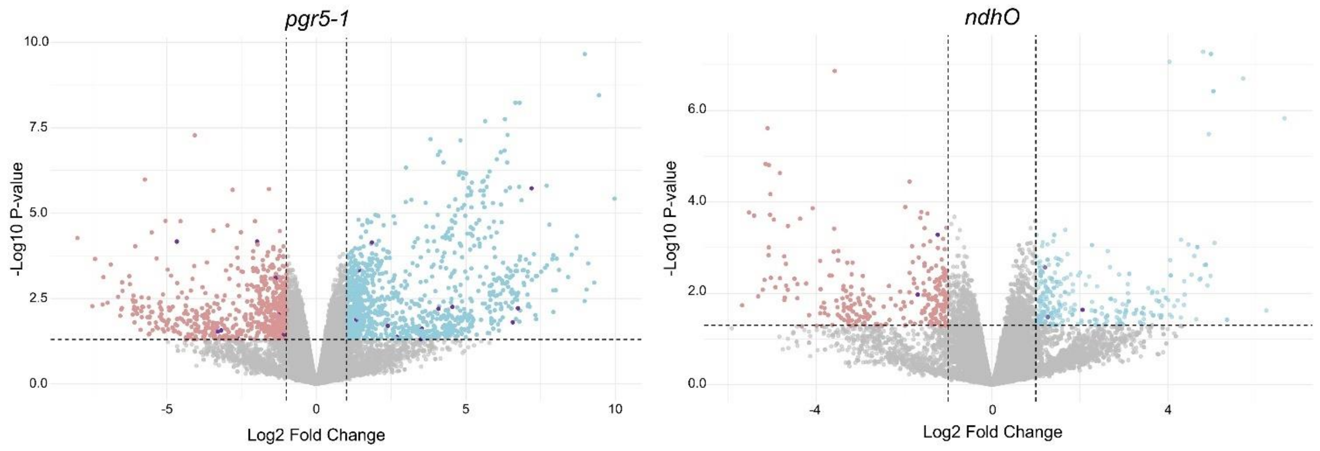


**Fig.S4.** Differential gene expression in *pgr5* and *ndhO S. viridis* compared to WT determined from RNAseq analysis. Upregulated genes are depicted in blue, while downregulated genes are depicted in red. Genes associated with Redox, ROS metabolism and oxidative stress are depicted in purple. PGR5-deficient plants demonstrated 561 upregulated genes and 282 downregulated genes compared to WT, while NDH-deficient plants demonstrated 114 upregulated genes and 127 downregulated genes compared to WT. NDH-deficient plants are reported in (Ermakova et al., 2024). ($n$ = 3 biological replicates, $p$ < 0.05, absolute $Log_2FC$ > 1)

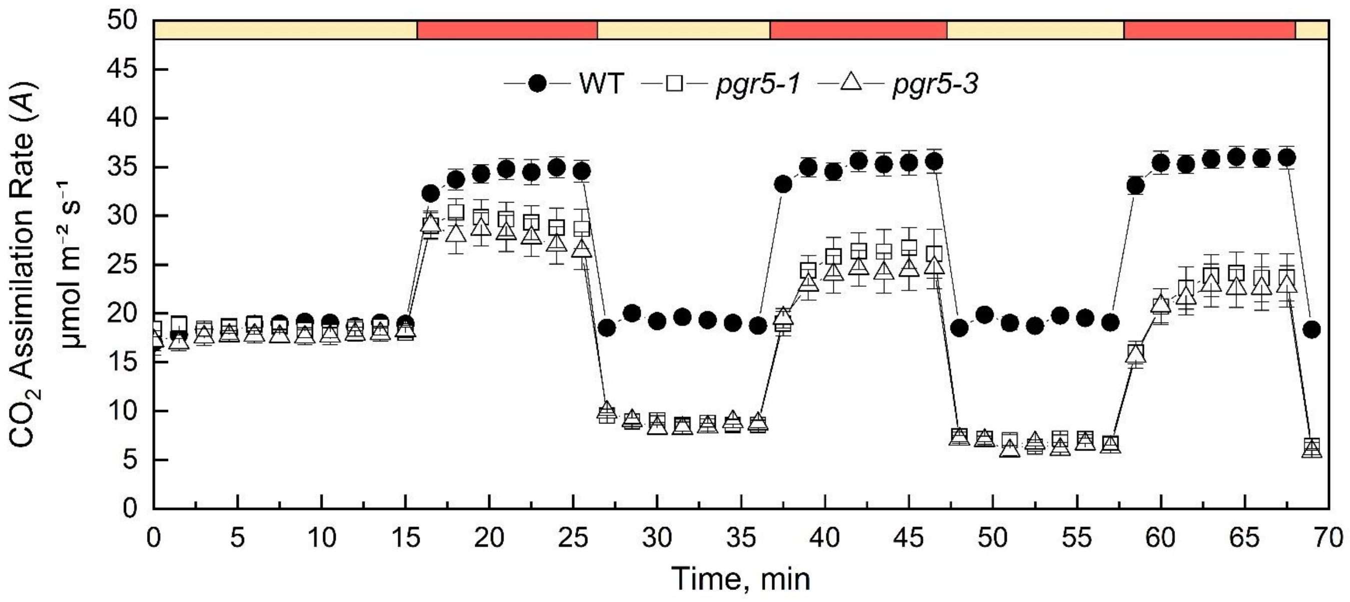


**Fig. S5**. Net $CO_2$ assimilation rate of WT *S.viridis* and PGR5-deficient plants (*pgr5-1* and *pgr5-3*) grown under constant daylight and subjected to fluctuating light sequence. Measurements were performed at ambient $CO_2$ partial pressure (400 µmol mol$^{-1}$). Coloured bars indicate the irradiance during the measurement: yellow, 400 µmol m$^{-2}$ s$^{-1}$; red, 2000 µmol m$^{-2}$ s$^{-1}$. Mean ± SE, $n$ = 5 biological replicates.
